# Poly dispersed acid-functionalized single walled carbon nanotubes target activated T and B cells to suppress GVHD in mouse model

**DOI:** 10.1101/2020.02.29.970996

**Authors:** Md. Babu Mia, Rajiv K. Saxena

## Abstract

Graft versus host disease (GVHD) results from hyper-activation of transplanted lymphocytes against the host antigens. Bone marrow transplantation in humans as well as some cases of blood transfusion and organ transplantation are associated with a strong GVH reaction resulting in GVHD that in many cases may be fatal. We had previously shown that poly-dispersed acid-functionalized single-walled carbon nanotubes (AF-SWCNTs) specifically target activated T and B lymphocytes and kill them. In the present study, efficacy of AF-SWCNTs to suppress the GVH reaction was tested in the mouse model. Acute GVHD was induced in mice by administering intravenously 30 or 60 million spleen cells from a parental strain (C57bl/6 mouse, MHC haplotype H-2^b^) to host (C57bl/6 × Balb/c) F1 mice (MHC haplotype H-2^b/d^) and waiting for 8-10 days. Chronic GVHD was similarly induced by administration of 30 million parent spleen cells to F1 mice and waiting for a period of 60 days. Our results demonstrate a marked decline in splenomegaly and recovery of spleen T (both CD4 and CD8) and B cells in GVHD mice treated with AF-SWCNTs. AF-SWCNTs treatment also limited T and B cell proliferation by restricting S-phage of cell cycle. Generation of anti-host cytotoxic T cells (CTLs) was also markedly suppressed by AF-SWCNT treatment of acute GVHD mice, and a significant reduction in the generation of anti-host antibodies could also be demonstrated. Taken together, our results suggest that the AF-SWCNTs can be considered as a potential therapeutic agent for treating GVHD.

## Introduction

Allogeneic Hematopoietic stem cell transplantation (HSCT) is the only curative therapy for a spectrum of hematological malignant and non-malignant disorders (Appelbaum, 2001). However, Graft-versus-Host disease (GVHD), a frequent, major, and fatal complication following HSCT, limits its therapeutic success (Ferrara and Deeg, 1991). In HSCT patients, acute GVHD (aGVHD) may result in 15-40 % mortality and up to 50 % morbidity (Choi et al., 2010, Ferrara and Deeg, 1991). Though GVHD occurs rarely in cases of solid organ transplants and blood transfusions to immune-incompetent hosts, but causes 90-100% mortality when it does occur (Chen et al., 2012, Rühl et al., 2009). Steroids are the drugs of choice for treating GVHD but there are no good alternative therapeutic approaches in cases where steroid therapy is ineffective (Martin et al., 2012). Multiple murine models have been developed to study GVHD (Schroeder and DiPersio, 2011). Administration of spleen cells of parent strain to F1 mice results in a potent GVH reaction since the immune system of the host F1 mouse ignores grafted parental spleen cells expressing self MHC, but the grafted spleen cells are free to raise a robust immune response to the MHC molecules inherited in F1 mouse from the other parental strain (van Bekkum and de VRIES, 1967, Gleichmann and Gleichmann, 1985, Moser et al., 1985). Various acute and chronic GVHD models have been used to study the disease mechanism or check the therapeutic effectivity of drugs. Developing the specific type of GVHD depends upon the activation of donor allogeneic CD4 & CD8 T cells (aGVHD) or CD4 T cells only in chronic GVHD (cGVHD) (Shearer and Levy, 1983, Hakim et al., 1991).

Nanomedicine is an emerging field that uses nanoparticles, including carbon nanotubes (CNTs), nano diamonds, gold nanoparticles, etc. for diagnostic and therapeutic purposes (Delogu et al., 2009). Within nanoparticles, carbon nanotubes (CNTs) with its unique physio-chemical structure (Niyogi et al., 2002), are finding wide applications in the fields of drug delivery (Flores et al., 2020, Bhaskar et al., 2010) vaccine delivery (Sacchetti et al., 2013), gene therapy (Cai et al., 2005) & tissue engineering (Aldinucci et al., 2013, Liu et al., 2009). Acid functionalization of single-walled CNTs significantly reduces their *in vivo* toxicity (Schipper et al., 2008) and enhances their interaction with immune cells (Saxena et al., 2007, Gazia and El-Magd, 2019, Gilmour et al., 2007). Previous studies have shown that the acid functionalized SWCNTs (AF-SWCNTs) modulate erythropoiesis (Bhardwaj and Saxena, 2015, Md and Saxena, 2018, Sachar and Saxena, 2011), downregulate protein and lipid antigen presentation (Rizvi et al., 2015, Kumari et al., 2012), alters epithelial tight junctions (Singh et al., 2020), reduce dendritic cells polarization capacity (Aldinucci et al., 2013), suppress activation of NK cells (Alam et al., 2016) and generation of alloimmune cytotoxic T cells (Alam et al., 2013, Dutt and Saxena, 2019). Interestingly, AF-SWCNTs preferentially target activated B and T cells and are cytotoxic to these cells (Dutt and Saxena, 2019, Dutt et al., 2019). In view of the general immunosuppressive effects of AF-SWCNTs and their relative affinity for activated T and B cells, they form a potential candidate for inducing immunosuppression and ablation of activated T and B lymphocytes *in vivo*.

In the present study, we have evaluated the effect of AF-SWCNTs in the mouse GVHD model. GVHD was induced in mice by injecting spleen cells from parental strain into F1 mice and effects of administration of AF-SWCNTs on the magnitude of the GVH reaction induced in this model was tested using various relevant parameters. We found that intravenous administration of AF-SWCNTs had a marked suppressive effect on the GVH reaction in this mouse model. Splenomegaly and high spleen cell recoveries of T and B cells characteristic of GVHD were significantly suppressed by AF-SWCNTs. A cytostatic effect of AF-SWCNTs on activated T and B cells is suggested by a significant decline in the proportions of T and B cells in S phase of cell cycle. Generation of anti-host cytotoxic T cells and antibodies were also significantly lower in AF-SWCNT treated GVHD mice. Taken together, our data suggest that AF-SWCNTs may be considered a viable candidate for further exploration as a treatment of GVDH.

## Methods and materials

### Animals

C57bl/6 (H-2^b^) and F1 mice (C57bl/6 × Balb/c F1) (8-12 weeks old, 20-25 g body weight) were obtained from Hylasco Biotechnology, Hyderabad. Animals were housed in pathogen-free, air-conditioned (25°C, 50% relative humidity) and 12 h light/dark cycle environment. Water and chow were provided ad libitum. All experimental protocols were conducted strictly in compliance with the guidelines notified by the Committee for Control and Supervision on Experiments on Animals (CPCSEA), Ministry of Environment and Forest (www.envfor.nic.in/divisions/awd/cpcsea_laboratory.pdf) and duly approved by the institutional animal ethics committee.

### Cells and reagents

Single walled carbon nanotubes (SWCNTs), N-hydroxysuccinimide (NHS), 1-ethyl-3-(3-dimethylaminopropyl) carbodiimide (EDAC), 2-(N-morpholino) ethane-sulfonic acid (MES) were purchased from Sigma Aldrich (St. Louis, MO). Centricon 3 kDa centrifugal filter device and 100 kDa dialysis tubing were ordered from Millipore (Billerica, MA). Alexafluor 633 hydrazide was from Molecular Probes (Carlsbad, CA). The P815 cells were purchased from ATCC (Virginia, USA). DMEM medium (with 4.5gms glucose, L-Glutamine, 3.7gms of sodium bicarbonate and sodium pyruvate / liter and 20μg/ml gentamycin), RPMI-1640 medium (with 2 mM glutamine, 1 mM sodium pyruvate, 4.5 g glucose/L, 10 mM HEPES, 1.5 g/L sodium bicarbonate and 20 lg/mL gentamycin) and fetal bovine serum (FBS) were purchased from Gibco (Carlsbad, CA). Purified anti-mouse CD16/32, fluorochrome tagged anti-mouse CD3, anti-mouse CD4, anti-mouse CD8, anti-mouse CD19, anti-mouse CD45, anti-mouse IgG / IgM antibodies were purchased from Biolegend (San Diego, CA). Carboxyfluoresceinsuccinimidyl ester (CFSE) was from eBiosciences (San Diego, CA). Propidium iodide and ribonuclease A (RNase A) were from Invitrogen (Carlsbad, CA).

### Acid functionalization of SWCNTs

Acid functionalization of SWCNTs was performed as described before (Alam et al., 2013). In short, SWCNTs (Sigma, Cat# 775535, > 95% carbon) (20mg) were suspended in 1:1 mixture of concentrated H_2_SO_4_ and HNO_3_ (20ml) and subjected to high pressure microwave (20 ± 2 psi, 50% of 900 W power, and temperature 135–150°C) for 3 min. Suspension was cooled, diluted 2X in water, dialyzed against deionized water, and lyophilized for further use. Zeta sizer analysis of AF-SWCNTs indicated an average particle size of 277.06 ± 3.37 d.nm and an average zeta potential of −47.02 ± 0.69 mV (Supplementary Figure 1 (A-B). BET surface area of SWCNTs was 799.399 M^2^/g and 240.741 M^2^/g for similar AF-SWCNTs measured by using flow BET nitrogen adsorption technique (Dutt et al., 2019). Transmission electron microscopic (TEM) examination revealed that acid functionalization has significantly reduced carbon nanotubes agglomeration without disturbing their basic structure (Supplementary Figure-1(C-D). To obtain Fluorescent tagged AF-SWCNTs (FAF-SWCNTs), AF-SWCNTs were covalently tagged with fluorochrome Alexa fluor 633 as described elsewhere (Alam et al., 2013).

### Developing GVHD models and AF-SWCNTs treatment

Three classical graft versus host disease (GVHD) models were used based upon previously established protocols (Schroeder and DiPersio, 2011) and used throughout the study. For acute GVHD-1 (aGVHD-1), spleen cells (30×10^6^, in 250 μl PBS) from female C57bl/6 mice were intravenously administered into female F1 mice and host mice sacrificed after 8 days. For acute GVHD-2 (aGVHD-2), spleen cells (60×10^6^, in 250 μl PBS) from female C57bl/6 mice were intravenously administered into female F1 mice and host mice sacrificed 10 days later. For chronic GVHD (cGVHD), spleen cells (30×10^6^, in 250 μl PBS) from female C57bl/6 mice were intravenously administered into female F1 mice and host mice sacrificed after 60 days. GVHD mice in each case were divided into control and AF-SWCNTs treated groups. AF-SWCNTs treated mice received three doses of AF-SWCNTs (50μg in 100 μl PBS) intravenously on 2^nd^, 4^th^ & 6^th^ days post administration of C57bl/6 mouse spleen cells into F1 mice, except aGVHD-2 mice that received one extra AF-SWCNTs dose on 8^th^ day. Control groups of mice received equal volume of PBS at same time points.

### In vivo uptake of FAF-SWCNTs by T and B cells

*In vivo* uptake of fluorescence tagged AF-SWCNTs (FAF-SWCNTs) was studied in aGVHD-2 mice. Four hours before sacrifice, control and AF-SWCNTs treated mice were administered 50μg of FAF-SWCNTs in 100μl of PBS through tail vein injection. Spleen cells from these mice were counterstained with CD3 and CD19 antibodies and uptake of FAF-SWCNTs in CD3^+^ T cells and CD19^+^ B cells was estimated by flow cytometry.

### In vitro cellular cytotoxicity assay

Generation of anti-H-2^d^ cytotoxic T cells was examined in aGVHD-2 mice using P815 mastocytoma (H-2^d^) as target cells. P815 cells were labeled with 10μM carboxyfluorescein succinimidyl ester (CFSE) for 10 minutes in ice, followed by 2 washings with PBS (Dutt et al., 2019). CFSE stains cells without any effect on their viability or functions. Spleen cells from control and AF-SWCNTs treated aGVHD-2 mice (10 × 10^6^) were mixed with CFSE labeled P815 cells (0.1 × 10^6^) in 100 μl of DMEM culture medium in 96 well U-bottom plates, that would result in an effector to target (E/T) ratio of 100:1. In control wells, P815 cells were incubated alone in equal volume of medium. Assay plates were centrifuged at 800 rpm for 5 minutes followed by incubation at 37°C for 8 hours. Cells in assay wells were re-suspended to get 100 μl of test cell suspension (TCS) in each assay well.

Separately, a calibrating preparation of CD45 stained spleen cells (CSCs) from control mice was kept ready. CSCs were prepared by using spleen cells from control F1 mice, depleted of erythrocytes by treatment with ACK (Ammonium-Chloride-Potassium) lysing buffer (Schmaltz et al., 2001), stained with leucocyte common antigen (anti CD45) antibody coupled to APC, washed and suspended at a concentration of 10^6^ cells/ml.

To 100 μl of TCS in each assay well, 100 μl of CSC suspension containing 0.1 × 10^6^ CSCs cells were added immediately before analysis of the samples on a flow cytometry so that the ratio of CFSE stained P815 cells to CD45 stained calibrating cells is 1.0 if all P815 cells survived. Thus, the P815 to CSC ratio in control wells (only P815 cells and no effector cells) would represent no specific killing of P815 cells except some background lysis. In test wells, where P815 cells were lysed by cytotoxic effector T lymphocytes (CTLs), the ratio of P815 to CSCs would decline. The ratio of P815 and CSCs was determined by flow cytometry for each assay well and percent lysis of P815 cells calculated by following formula:

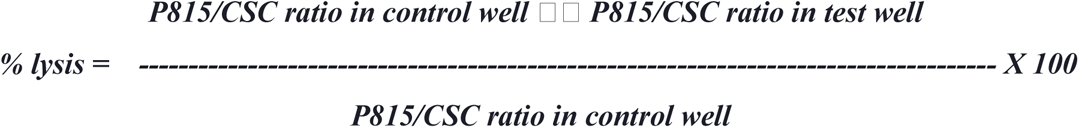

### Assessment of auto-antibodies in GVHD mice

A C57bl/6 vs. (C57bl/6 × Balb/c) F1 GVHD reaction may generate anti-host (H-2^b^) antibodies made by grafted and activated B cells that would recognize host antigens as non-self. To assess the levels of anti-host autoantibodies, sera from GVHD mice were used to stain thymocytes from F1 mice. cGVHD mice were sacrificed at 60^th^ days, and blood samples were collected to isolate serum. Thymocytes (0.1 × 10^6^) derived from control F1 mice were incubated with cGVHD mouse sera at 1: 10 dilutions for 30 minutes in ice and washed twice with PBS. Cells were further stained with anti-mouse IgG / IgM secondary antibodies to reveal thymocytes with membrane bound antibodies. Thymocytes were chosen for this purpose as they do not have membrane Ig receptor that would otherwise cause false positives by reaction with the secondary antibodies in this assay system. Proportion of stained thymocytes using different sera from GVHD mice would indicate the relative levels of auto-antibodies.

### Flow cytometric assay

Spleen cells/ thymocytes to be analyzed were incubated first with anti-mouse CD 16/32 for 30 minutes in ice to block Fc receptors. Cells were then stained with specific fluorochrome tagged anti-mouse antibodies (CD3, CD4, CD8, and CD19) as per the instructions for using the antibodies and analyzed on flow cytometer. For studying cell cycle in T and B cells, stained cells were fixed overnight using 70% ethanol at 4°C in the dark. Ethanol washed cells were treated with Ribonuclease A (50μg/ml, 1h at 37°C) and stained with Propidium Iodide (5μg/ml) for 30 minutes in ice. 10 to 100 thousand events were accumulated for flow cytometric analysis. Flow cytometric analysis was performed using FACSuite, FlowJo v10 & Modfit v3.1 software.

### Statistical analysis

Each GVHD experiments were repeated three times with minimum three mouse in each group. Data analysis was performed using Sigma Plot software (Systat Software, San Jose, CA) and presented as Mean ± SEM. Differences between groups were calculated using student t-test or ANOVA. P value < 0.05 was accounted as significant (P = <0.05 *, <0.01 **, <0.005 ***, <0.001 ****).

## Results

### AF-SWCNTs suppress GVHD associated splenomegaly and restrict clonal expansion of allo-reactive cells

Administration of parental strain spleen cells into F1 mouse results in a GVH reaction since the grafted cells react against the host but host’s immune system ignores the grafted cells as they express syngeneic histocompatibility antigens. Robust proliferation of activated host lymphocytes in mice undergoing GVH reaction results in splenomegaly. Spleens isolated from mice undergoing GVH reaction (all three protocols i.e. aGVHD-1, aGVHD-2, cGVHD) with or without AF-SWCNTs treatment are shown in Figure 1A where splenomegaly is apparent, especially in the aGVHD-2 model. Statistically significant increase in the spleen weights was observed in aGVHD-1 as well as aGVHD-2 mice (Figure 1B) but not for cGVHD mice that were sacrificed 60 days after the induction of GVH reaction (Figure 1B). In case of aGVHD-2 mice, a statistically significant decline in spleen weight was seen in AF-SWCNTs treated mice (Figure 1B). Figure 1C shows that the total spleen cell recoveries were markedly elevated in all models of GVHD as compared to control non-GVHD mice. A significant decline in spleen cell recovery was clearly observed as a result of treatment with AF-SWCNTs in all three models of GVHD (aGVHD-1 23%↓, aGVHD-2 61 %↓, cGVHD 40 %↓) (Figure 1C). These results suggest that splenomegaly and spleen cell recoveries in mice undergoing GVHD are suppressed significantly by treatment with AF-SWCNTs.

**Figure 1:**
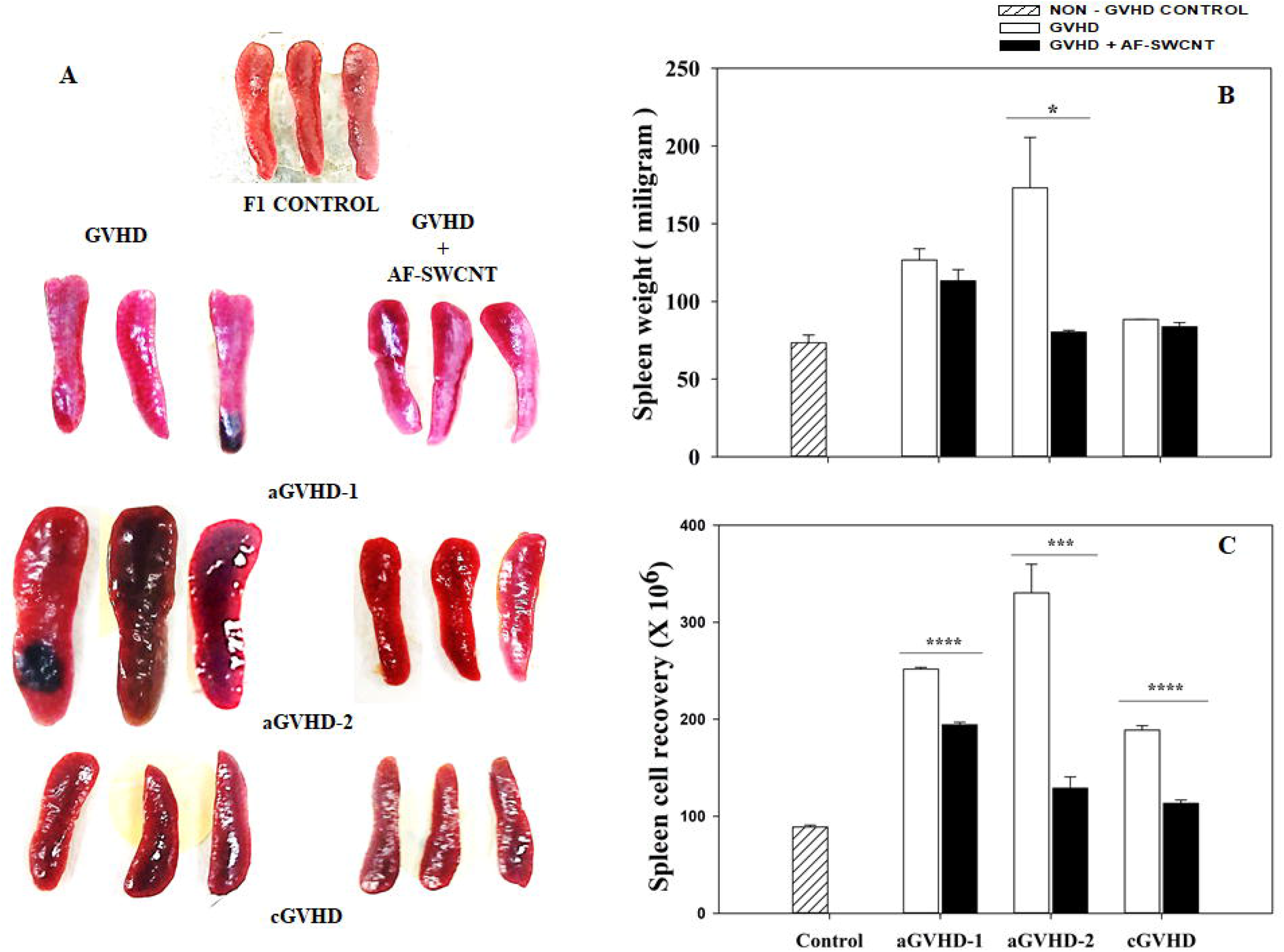
Decrease in spleen size and cell recovery in GVDH mice treated with AF-SWCNTs. Spleen cells from C57bl/6 mice were injected intravenously in F1 mice as described in methods. Treated groups were administered 3 or 4 doses of AF-SWCNTs (50 μg per dose) as described in methods. Panel A shows the photographs of spleens from control, GVHD and GVHD+AF-SWCNT groups of mice. Panel B shows the data on spleen weights and Panel C shows the data on total spleen cell recoveries in different groups of mice. All results represent Mean ± SEM of replicative experiments of each individual group. (**p< 0.01, ****p<0.001)

### AF-SWCNTs suppress the recovery of T cells and B cells from spleens of GVHD mice

GVHD is caused by proliferating T and B lymphocytes from grafted spleen cells within the host mouse. Proportions as well as absolute recoveries of CD3^+^ T cells and CD19^+^ B cells were enumerated by flow cytometry in all three models of GVHD. Results in Figure 2 show that the proportions of CD3^+^ T cells (Figure 2A) as well as CD19^+^ B cells (Figure 2B) increased markedly in GVDH mice and fell significantly in GVHD mice treated with AF-SWCNTs. Absolute recoveries of both T and B cells were 4 to 10-fold higher in GVHD mice as compared to control non-GVHD mice and these elevated recoveries fell significantly as a result of AF-SWCNT treatment (30 to 70%) in all models of GVHD (Figure 2 C, D). Changes in the proportions and absolute recoveries of CD4 and CD8 subsets of T cells in GVHD control and AF-SWCNT treated groups were also assessed and are shown in Figure 3. Both proportions and absolute recoveries of T cell subsets were also significantly reduced in GVHD mice treated with AF-SWCNTs.

**Figure 2:**
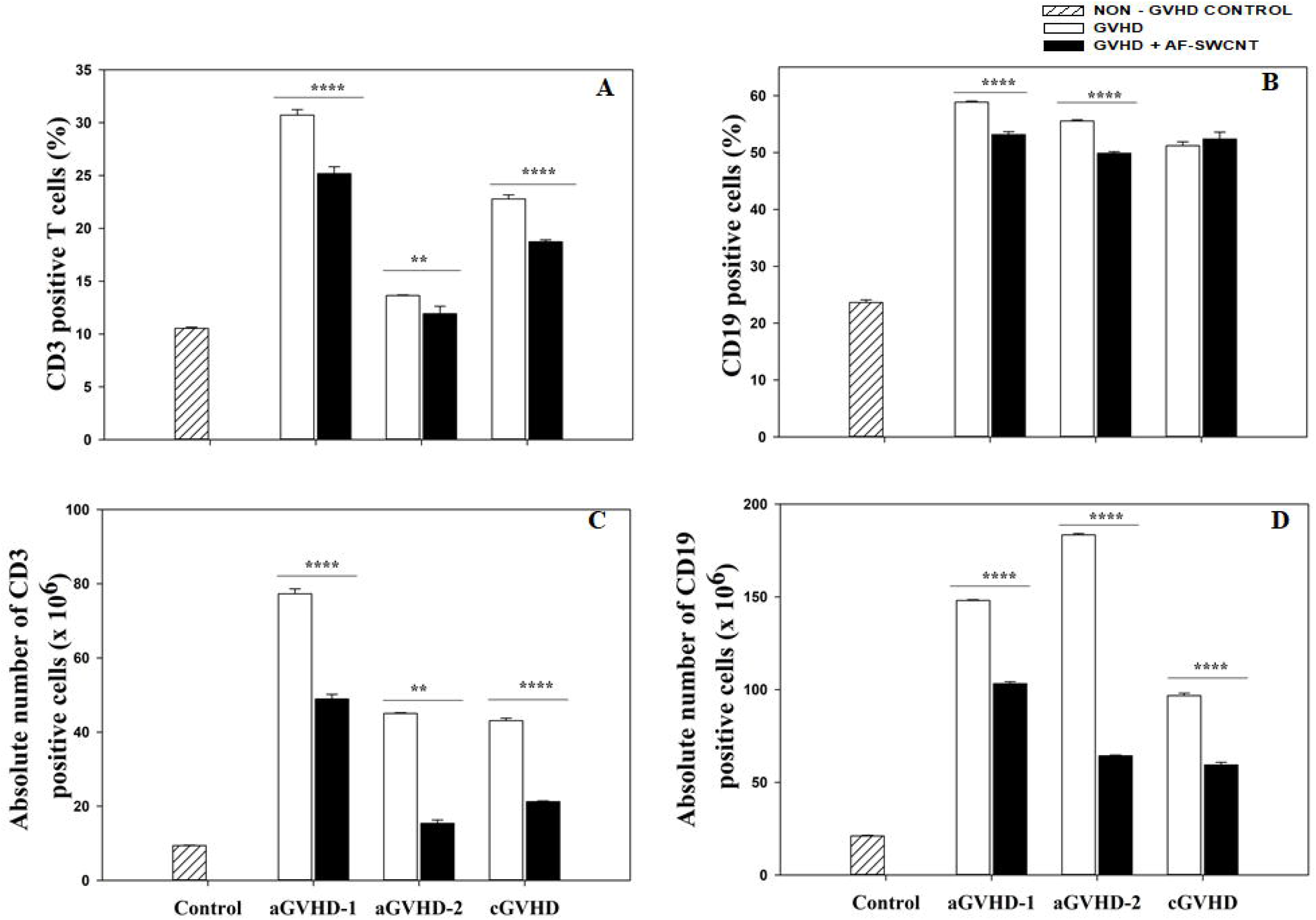
Effect of AF-SWCNT treatment on recoveries of T and B cells from GVHD mice. Control group of mice, GVDH mouse groups of three types as explained in methods and GVDH+AF-SWCNT group of mice were sacrificed and recovery of CD3^+^ T cells and CD19^+^ B cells were determined. Percentages of CD3^+^ T and CD19^+^ B cells in spleen cell suspensions determined by flow cytometry are shown in panels A and B. Absolute CD3^+^ T and CD19^+^ B cells recovery per spleen are given in panels C and D. All results represent Mean ± SEM of replicative experiments of each individual group. (**p< 0.01, ****p<0.001)

**Figure 3:**
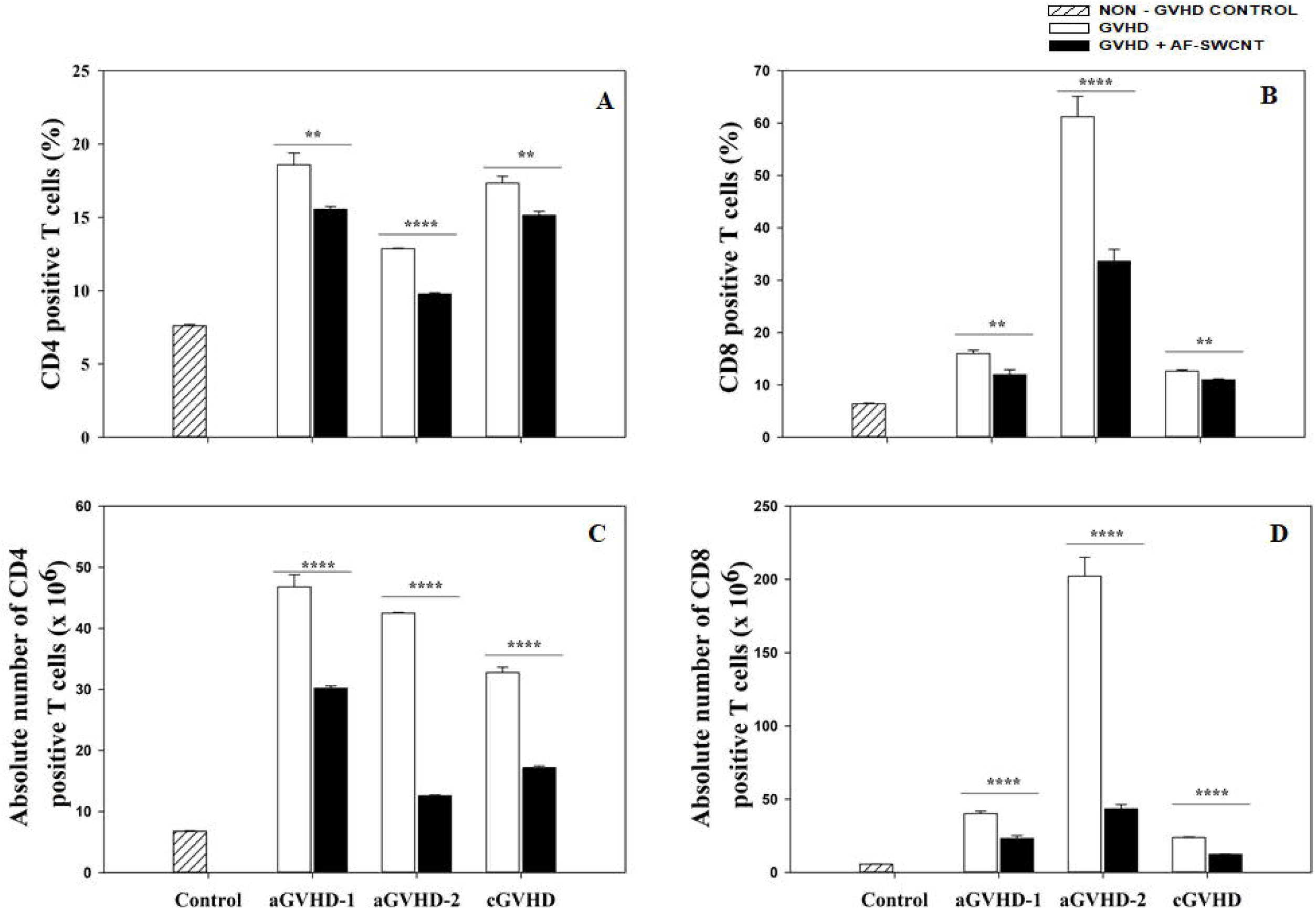
Recoveries of CD4 and CD8 T cells from spleens of control, GVHD and GVHD+AF-SWCNT groups of mice. Control group of mice, GVDH mouse groups of three types as explained in methods and GVDH+AF-SWCNT group of mice were sacrificed and recovery of CD4^+^ and CD8^+^ T cells were determined. Percentages of CD4^+^ and CD8^+^ T cells in spleen cell suspensions were determined by flow cytometry (panels A and B). Absolute recoveries per spleen of CD4^+^ and CD8^+^ T cells are given in panels C and D. All results represent Mean ± SEM of replicative experiments of each individual group. (**p< 0.01, ****p<0.001)

### In vivo uptake of FAF-SWCNTs by activated T and B cells in GVHD mice

Reduction of the recoveries of T and B cells in AF-SWCNTs treated GVHD mice could result from a reduction in the proliferation of these cells or killing of these cells by AF-SWCNTs. We have previously shown that the uptake of AF-SWCNTs in activated T and B cells is markedly higher as compared to resting control T and B cells (Dutt and Saxena, 2019, Dutt et al., 2019). A reduction in the uptake of fluorescence tagged AF-SWCNTs (FAF-SWCNTs) in AF-SWCNT treated GVHD mice would be a pointer to a reduction of activation status of T and B cells. Relative uptake of AF-SWCNTs by T and B cells in GVHD control and AF-SWCNT treated mice was assessed by using fluorescence tagged AF-SWCNTs (FAF-SWCNTs). FAF-SWCNTs were administered intravenously in aGVHD-2 mice (GVHD and AF-SWCNTs treated GVHD groups), four hours before the sacrifice. Spleen cells derived from these mice were stained with anti-mouse CD3 and CD19 antibodies followed by flow cytometric analysis. FAF-SWCNT stained cells within CD3^+^ T cell and CD19^+^ B cell populations were enumerated. As per the data shown in Figure 4, 22.92 ± 0.36 % CD3^+^ T cells from control aGVHD-2 mice were FAF-SWCNT positive (MFI = 244.4 ± 5.88), which was substantially reduced to 3.94 ± 0.13 % T cells (MFI = 153.33 ± 0.54) in AF-SWCNTs treated mice (Figure-4 A,B). These results indicate that the uptake of AF-SWCNTs by T cells in aGVHD-2 mice was significantly reduced (82.81%) upon AF-SWCNTs treatment. Similarly, FAF-SWCNTs uptake in CD19^+^ B cells was also significantly lower [7.47 ± 0.11 % (MFI =162.67 ± 0.25) in control aGVHD-2 mice to 1.76 ± 0.06 % (MFI = 140.67 ± 1.96) in AF-SWCNTs treatment group] after AF-SWCNTs treatment (Figure-4 C, D). These results indicate that the numbers of T and B cells that internalized FAF-SWCNTs were significantly reduced in AF-SWCNTs treated aGVHD-2 mice.

**Figure 4:**
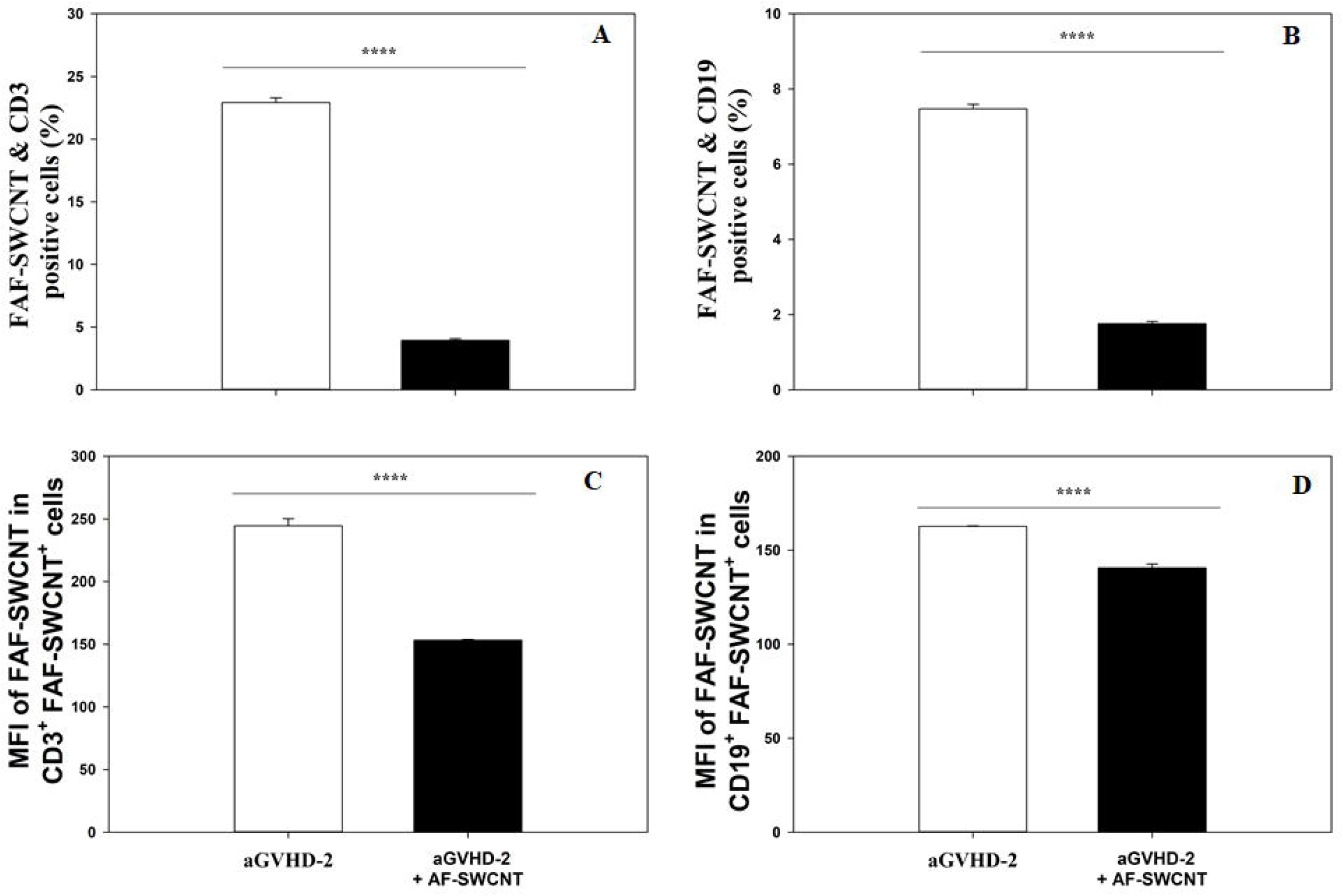
*In vivo* uptake of FAF-SWCNTs by T and B cells. Spleen cells from C57bl/6 mice were injected intravenously in F1 mice as described in methods. Treated groups were administered 3 or 4 doses of AF-SWCNTs (50 μg per dose) as described in methods. FAF-SWCNTs (50 μg in 100 μl PBS) was injected four hours before the sacrifice of all mice. All cell preparations were stained for T (CD3) and B (CD19) cells and cells with FAF-SWCNTs determined by flow cytometry for each cell preparation. Panels A and B show the percentage of FAF-SWCNT positive T and B cells respectively. Panels C and D shows the data on mean fluorescence intensity (MFI) of the FAF-SWCNT stain. All results represent Mean ± SEM of replicative experiments of each individual group. (**p< 0.01, ****p<0.001)

### Reduction of S phase of cell cycle in T and B cells from AF-SWCNTs treated aGVHD-2 mice

In order to see if AF-SWCNT treatment had an effect on the cell cycle of proliferating T and B cells in GVHD mice, spleen cells from aGVHD-2 control and AF-SWCNT treated mice were stained for CD3 and CD19 cells and further stained with propidium iodide to gate G0/G1, S, G2/M phases of the cell cycle as described in the methods & materials section. Results in Figure-5A show that a significant reduction in S phase occurred in T cell population (from 25.81 ± 2.47% to 16.51 ± 1.09%) as a result of AF-SWCNTs treatment (Figure 5A). Similarly, AF-SWCNTs treatment reduces CD19^+^ B cells in S-phase from 4.97 ± 0.11% to 0.71 ± 0.30% (Figure-5B).

**Figure 5:**
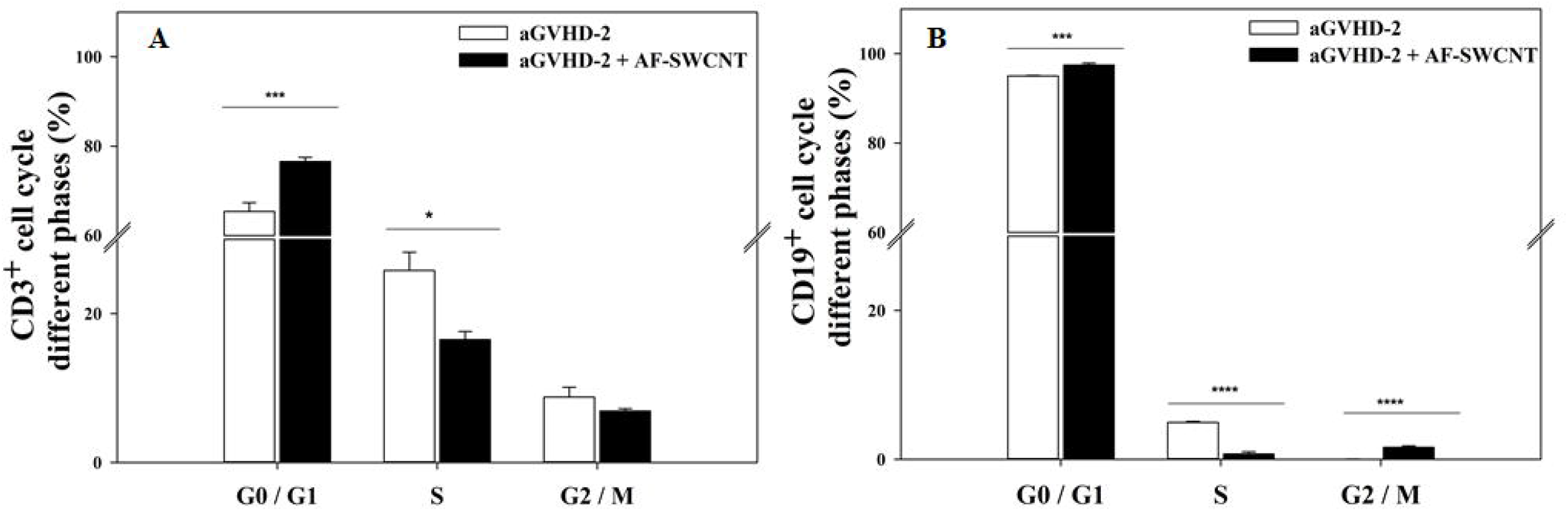
Effect of AF-SWCNT treatment on cell cycling patterns of T and B cells in GVHD and GVHD+AF-SWCNT groups of mice. aGVHD-2 mice with or without AF-SWCNT treatment were sacrificed and spleen cells were stained with mouse CD3 and CD19 antibodies followed by PI stain as described in methods. Percentage of cells in different stages of cell cycle within T cells (panel A) and B cells (panel B) populations was determined by flow cytometry. All results represent Mean ± SEM of replicative experiments of each individual group. (**p< 0.01, ****p<0.001)

### AF-SWCNT reduces the generation of allo-reactive cytotoxic T lymphocytes (CTLs) in GVHD mice

GVH reaction results in activation of cytotoxic T lymphocytes (CTLs) against cells bearing host histocompatibility antigens. In our system [b anti (b × d) F1], this cytotoxicity would be directed against H-2^d^ bearing cells. If AF-SWCNTs treatment suppressed the GVH reaction, it was important to see if the generation of anti-H-2^d^ allo-immune CTL response was suppressed in GVHD mice treated with AF-SWCNTs. For measuring the anti-H-2^d^ CTL response, we used P815 target cells that express H-2^d^. P815 cells were labelled with a fluorescent membrane binding dye (CFSE) and incubated with spleen cells containing CTLs for 8 hours. Percentage of P815 that were killed by CTLs was estimated by a flow cytometric assay explained in materials and methods. Results in Figure 6 show that about 10.36 ± 1.30 % of the P815 cells incubated with spleen cells from control non-GVH mice were killed that represented background killing. For spleen cells from aGVHD-2 mice, P815 cell killing went up to 37.06 ± 0.65 %. Further, in AF-SWCNTs treated aGVHD-2 mice, the P815 killing came down to 13.76 ± 0.70 %. These results clearly show that the generation of allo-immune CTLs in GVH is significantly suppressed by treatment with AF-SWCNTs.

**Figure 6:**
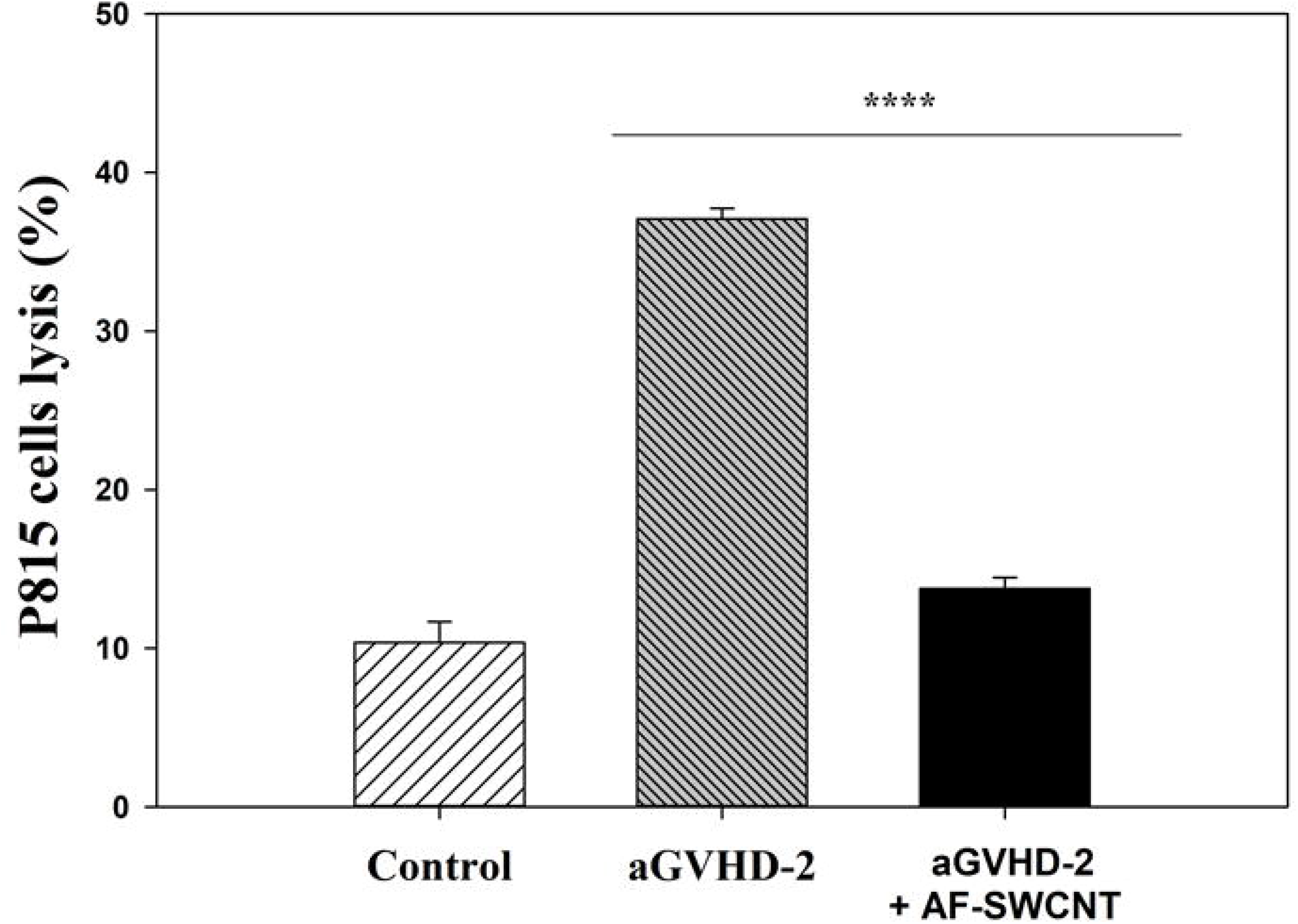
Anti-P815 cytotoxic activity in spleen cells derived from GVHD mice with or without AF-SWCNT treatment. Anti-P815 cytotoxic activity of spleen cells from non-GVHD control mice and GVHD mice with or without AF-SWCNT treatment was determined as described in methods. All results represent Mean ± SEM of replicative experiments of each individual group. (**p< 0.01, ****p<0.001)

### Suppression of auto-antibody response in chronic GVHD mice treated with AF-SWCNTs

Besides CTL response in GVH reaction, auto antibodies are also produced against the host cells (Shearer and Levy, 1983). Effect of AF-SWCNTs was therefore also examined on the production of auto-immune antibodies in GVHD mice. Since antibody response takes longer time, auto-antibody levels were assessed in sera from cGVHD mice with or without treatment with AF-SWCNTs. Auto-antibody response was tested using thymocytes from control F1 host mice. Thymocytes were treated with antisera at 1:10 dilution, and membrane bound autoantibodies on thymocytes assessed by staining with fluorochrome tagged secondary antibodies against mouse immunoglobulin, as described in materials and methods. Control thymocytes express no membrane immunoglobulin of their own and thus will not get stained with the secondary antibody reagent alone. Results in Figure 7 show that treatment with control mouse serum did not stain thymocytes. There was a significant staining of thymocytes when cGVHD serum was used (20.30 ± 0.53 %) and this stain was lower by about 15 % (p <0.007) when serum from AF-SWCNTs treated cGVHD mice was used. These results suggest that treatment with AF-SWCNTs may lower the production of autoantibodies in a GVH reaction.

**Figure 7:**
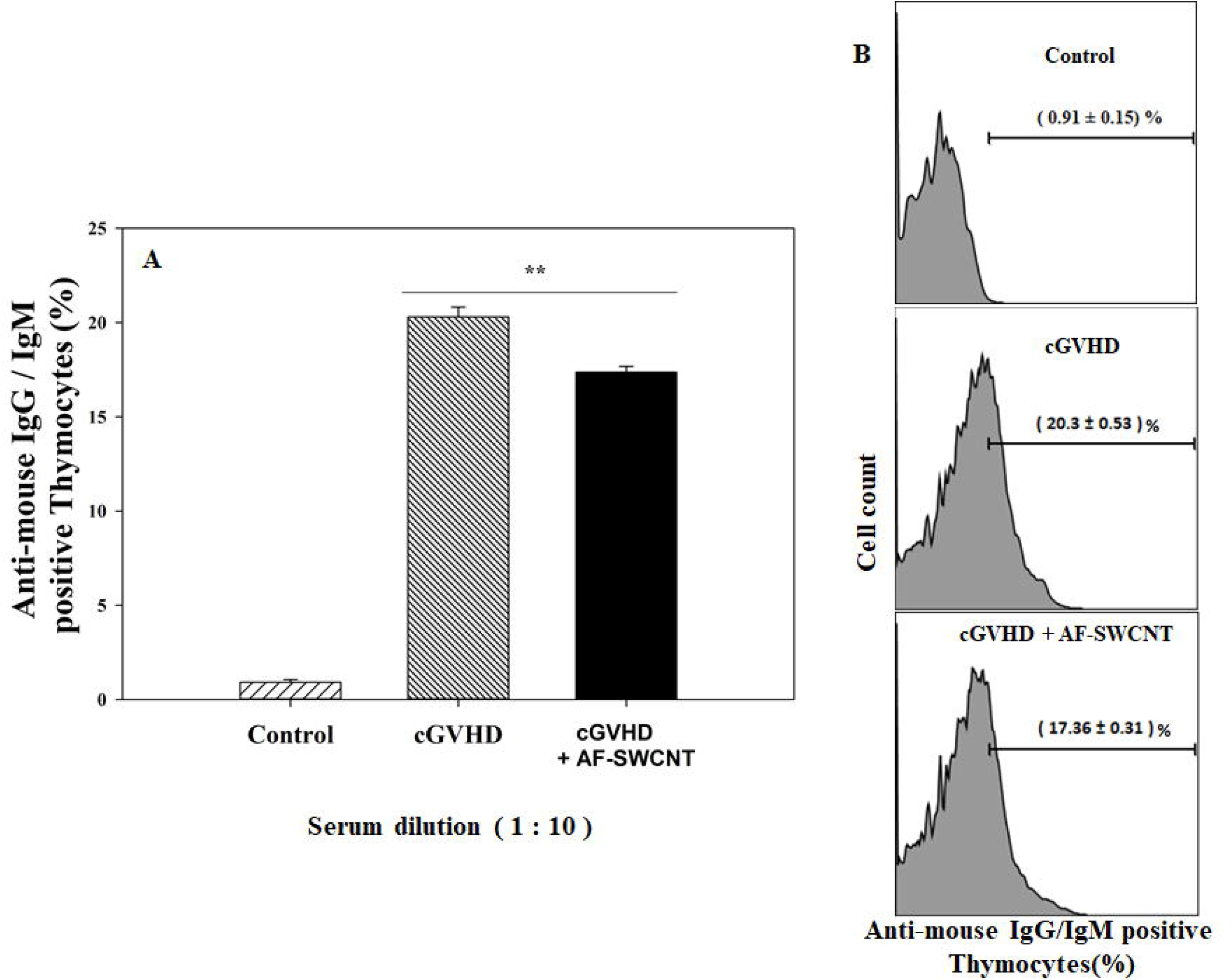
Anti-host antibody response in GVHD mice with or without AF-SWCNT treatment. Sera were collected from non-GVHD control mice and GVHD mice with or without AF-SWCNT treatment and presence of anti-host thymocyte antibodies determined as described in methods. All results represent Mean ± SEM of replicative experiments of each individual group. (**p< 0.01, ****p<0.001)

## Discussion

Work from our lab and others has shown the interaction of carbon nanotubes with resting and activated T (Alam et al., 2013, Dutt and Saxena, 2019, Bottini et al., 2006, Kim et al., 2016, Mitchell et al., 2009) and B cells (Dutt et al., 2019, Kim et al., 2016). In essence, activated T and B cells show enhanced uptake of acid functionalized single walled carbon nanotubes (AF-SWCNTs) as compared to resting T and B cells. Enhanced uptake of AF-SWCNTs may also lead to killing of activated lymphocytes (Kermanizadeh et al., 2016, Dutt and Saxena, 2019, Dutt et al., 2019, Alam et al., 2013). This finding opens up the possibility of using AF-SWCNTs to specifically kill activated T and B cells in clinical situations where deletion of activated populations of lymphocytes may be clinically helpful. This proposition has however not yet been examined experimentally. In the present study we have examined the efficacy of AF-SWCNTs to suppress the graft versus host reaction *in vivo*. Graft versus host disease (GVHD) is a frequent and fatal complication that arises following allogeneic bone marrow transplantations in humans (Appelbaum, 2001, Ferrara and Deeg, 1991, Choi et al., 2010). Steroids are used to suppress the GVHD but if that is ineffective, there are no other proven second line of therapies for steroid non-responders (Martin et al., 2012). GVH reaction can be induced in mice by injecting parent spleen cells to F1 mice (Schroeder and DiPersio, 2011). Parent to F1 GVHD mouse model mimics the human GVHD and presents an opportunity to test the efficacy of AF-SWCNTs in suppressing the GVH reaction.

Based upon previously established techniques (Gleichmann and Gleichmann, 1985, Moser et al., 1985), we used three types of protocols to induce GVHD in mice, explained in materials and methods. In all these protocols, spleen cells from C57bl/6 (MHC haplotype H-2^b^) were injected intravenously into (C57bl/6 × Balb/c) F1 mice (MHC haplotype H-2^b/d^). Nguyen et al., (2007) have demonstrated by *in vivo* bioluminescence study that the transplanted parent cells localize in spleen within 24 hours of administration. Spleen is therefore a major site where the grafted cells start a robust immune response to host histocompatibility antigens resulting in proliferation of activated T and B cells and a consequent increase in spleen size (splenomegaly). We found significant splenomegaly in the two models of aGVHD but the increases spleen weight was most pronounced in aGVHD-2. Significant splenomegaly was not seen in cGVHD. Nonetheless, the total cell recoveries from spleen was highly elevated in all three models of GVHD and treatment with AF-SWCNTs resulted in a significant decline in spleen size as well as spleen cell recovery in all cases. Both T and B cells get activated and proliferate in a GVH reaction (Schroeder and DiPersio, 2011, Rus et al., 1995). We found in general an increase in the proportions of T and B cells in spleens of GVHD mice, but the effect was especially pronounced in total spleen cell recoveries where increases ranging from 4 to 8-fold were observed. Spleen cell recoveries rather than proportion of cellular populations should be more significant in this case since proportions would depend upon relative changes in number of different spleen cell populations, whereas total recoveries would be a pointer to actual activation and proliferation of individual cell populations. Within the T cell population, recovery of spleen CD4 T cells were found to be 6 to 9-fold higher than control spleens in acute as well as chronic GVHD. Interestingly, maximum increase of the recovery of CD8 T cells was seen in aGVHD-2 mice (almost a 40-fold increase) though less pronounced yet significant increases were seen in aGVHD-1 and cGVHD mice. In all cases however, a substantial fall in CD4 and CD8 spleen recoveries occurred in GVHD mice treated with AF-SWCNTs. Further experiments were done to understand the mechanism of decline in T and B cell recoveries in response to AF-SWCNT treatment and since the most pronounced changes were observed in aGVHD-2 model, this was used for subsequent studies.

In the aGVHD-2 model we found that the fraction of T and B cell populations that internalize fluorescence tagged AF-SWCNTs (FAF-SWCNTs) *in vivo* fell markedly (80 to 90% decline) in AF-SWCNT treated aGVHD-2 mice. Average per T or B cell uptake of FAF-SWCNTs indicated by mean fluorescence intensities also fell by 20 to 40%. Since activated T and B cells internalize substantially high amounts of AF-SWCNTs (Dutt and Saxena, 2019, Dutt et al., 2019) the reduction we observed in FAF-SWCNT uptake by T and B cell populations in GVHD mice treated with AF-SWCNTs may be a reflection of steep decline in the numbers of activated T and B cells. Decline in T and B cell recoveries in AF-SWCNT treated GVHD mice could be due to blocking of cell division or cytotoxic effect of internalized AF-SWCNTs, or both. A significant decline in the proportion of T and B cells in S phase of cell cycle in AF-SWCNT treated aGVHD-2 mice suggests that the AF-SWCNT treatment had cytostatic effect. Direct killing of activated T and B cells by AF-SWCNTs also appear likely in view of a substantial drop in T and B cell recoveries, but further experimental evidence will be needed to confirm this proposition *in vivo*.

GVH disease is generally fatal and both cytotoxic T cells as well activated B cells producing anti-host antibodies may contribute to the fatality (Shearer and Levy, 1983, Shimabukuro-Vornhagen et al., 2009). It was therefore important to determine if the generation of anti-host CTLs and antibodies were also suppressed in AF-SWCNT treated GVHD mice. In the GVHD mouse model we used, generation of anti H-2^d^ CTLs is expected since the grafted C57bl/6 (H-2^b^) T cells would react to H-2^d^ expression on F1 host cell. To assess the anti H-2^d^ CTL activity, we used P815 (H-2^d^) target cells that are often used as CTL targets (Alam et al., 2013, Saxena et al., 1982). We also developed a flow cytometric CTL assay using fluorescence tagged P815 cells as targets and enumerating the surviving P815 cells in comparison to a fixed concentration of CD45 stained control F1 spleen cells used as a calibrator as described in methods. We found a substantial decline in the generation of anti-H-2^d^ CTLs (65% decline), that provides another rationale of a possible use of AF-SWCNTs for treatment of GVHD.

Chronic GVHD results in up to 50% morbidity after HSCT (Choi et al., 2010, Ferrara and Deeg, 1991), which may be caused by auto-antibody generated by grafted B cells (Morris et al., 1990, Kuzmina et al., 2015). Therefore, it was tempting to evaluate the AF-SWCNTs effect on auto-antibody formation in cGVHD. Since 8 to 10 days are not sufficient for the antibody response, cGVHD model was used for this purpose. Production of auto-antibodies was marginally, though significantly lower in AF-SWCNTs treated cGVHD mice. It should be noted that even in cGVHD model only three doses of AF-SWCNTs were given within first 8 days of the 60 days long cGVHD protocol. It is possible that a more sustained treatment with AF-SWCNTs might have further reduced the auto antibody production, though that will require further testing.

Taken together, our study demonstrates in the murine GVHD model that treatment with AF-SWCNTs may considerably reduce the vigor of the GVH reaction that otherwise kills the host. These results may lead to further studies aimed at testing the validity of the use of AF-SWCNTs as an alternative mode of treatment of human GVHD.

---

**Figure S1: Acid functionalized single walled carbon nanotubes (AF-SWCNTs) characterization.** Panel A shows the mean particle size distribution on a zeta sizer, and Panel B shows the mean zeta potential of AF-SWCNTs on zeta sizer. Panel C and D shows the Transmission Electron Microscopy (TEM) images of SWCNTs and AF-SWCNTs at 30000X magnification.

