## Supplementary figures and images for "Poly dispersed acid-functionalized single walled carbon nanotubes target activated T and B cells to suppress GVHD in mouse model"

### Supplementary data 1

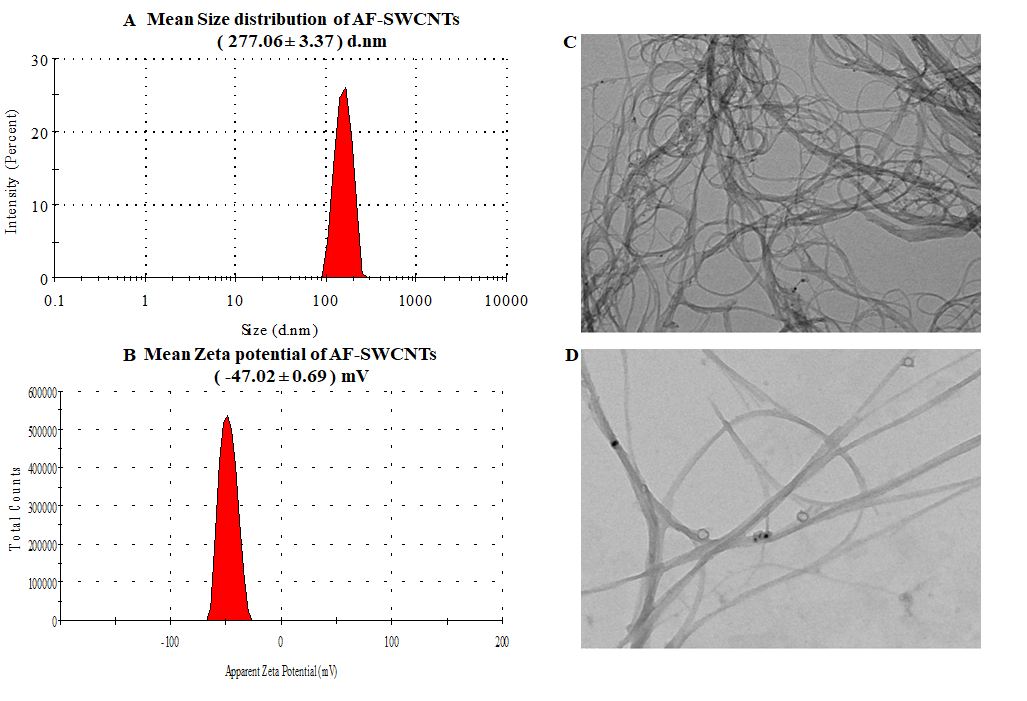
